# Mixotrophic microalgal mixed cultures for cheese whey valorization

**DOI:** 10.1101/2023.10.24.563819

**Authors:** Maria Paula Giulianetti de Almeida, Camille Mondini, Guillaume Bruant, Julien Tremblay, Gustavo Mockaitis, David G. Weissbrodt

**Affiliations:** Interinstitutional Graduate Program in Bioenergy (USP/UNICAMP/UNESP) – 330 Cora Coralina Street, Campinas/SP, CEP 13.083-896, Brazil; Interdisciplinary Research Group on Biotechnology Applied to the Agriculture and the Environment, School of Agricultural Engineering, University of Campinas (GBMA/FEAGRI/UNICAMP), Campinas, SP, Brazil; Department of Biotechnology, Delft University of Technology, Delft, The Netherlands; National Research Council Canada, Energy, Mining and Environment research centre, Genomics and microbiomes group, 6100 Royalmount Avenue, H4P 2R2, Montreal, QC, Canada; Department of Biotechnology and Food Science, Norwegian University of Science and Technology, N-7491 Trondheim, Norway

**Keywords:** Microalgae, phycoremediation, biomass contamination, anaerobic coupled processes, cheese whey, photoorganoheterotrophy

## Abstract

Microalgae cultivation, and phycoremediation, can be a polishing step in wastewater treatment. Depending on the stream utilized for microalgal cultivation, biomass can be contaminated with considerable quantities of heavy metals and xenobiotics. Given the economic value of microalgae bioproducts, we suggest coupling anaerobic fermentation with microalgae mixotrophic growth. Cheese whey, a product from cheese production, has a 2022 forecast production of 160.7 million m^3^ year^−1^ in which about 66.5 million m^3^ y^−1^ is used as animal feed, fertilizers or illegally discharged causing eutrophication. Anaerobic fermentation of cheese whey produces volatile fatty acids (VFAs) such as acetate which serves as an organic carbon source for photoorganoheterotrophic microalgal biomass growth. Our work selected three organic sources derived from cheese whey: 40% demineralized whey powder (WPC40), lactose, and acetate. In photolitoautotrophic conditions, green phototrophic growth was successful. In batch tests, acetate was the best organic carbon source among photoorganoheterotrophs with a higher yield of 1.15 mg VSS mg Carbon^−1^ (C) in anaerobic conditions. Also, acetate uptake was thought to be via the glyoxylate cycle. When upscaling the experiment in a chemostat, a lower dilution rate of 0.17 d^−1^ was more suitable for green photoorganoheterotrophs selection, as they were not washed out in the process. These findings show that acetate uptake by microalgae in mixotrophic regimes must be better understood as well as reinforce the advantages of coupling microalgal biomass growth with cheese whey acidogenic fermentation, avoiding contaminations as in phycoremediation processes and fully valorizing cheese whey.

**Graphical abstract:** 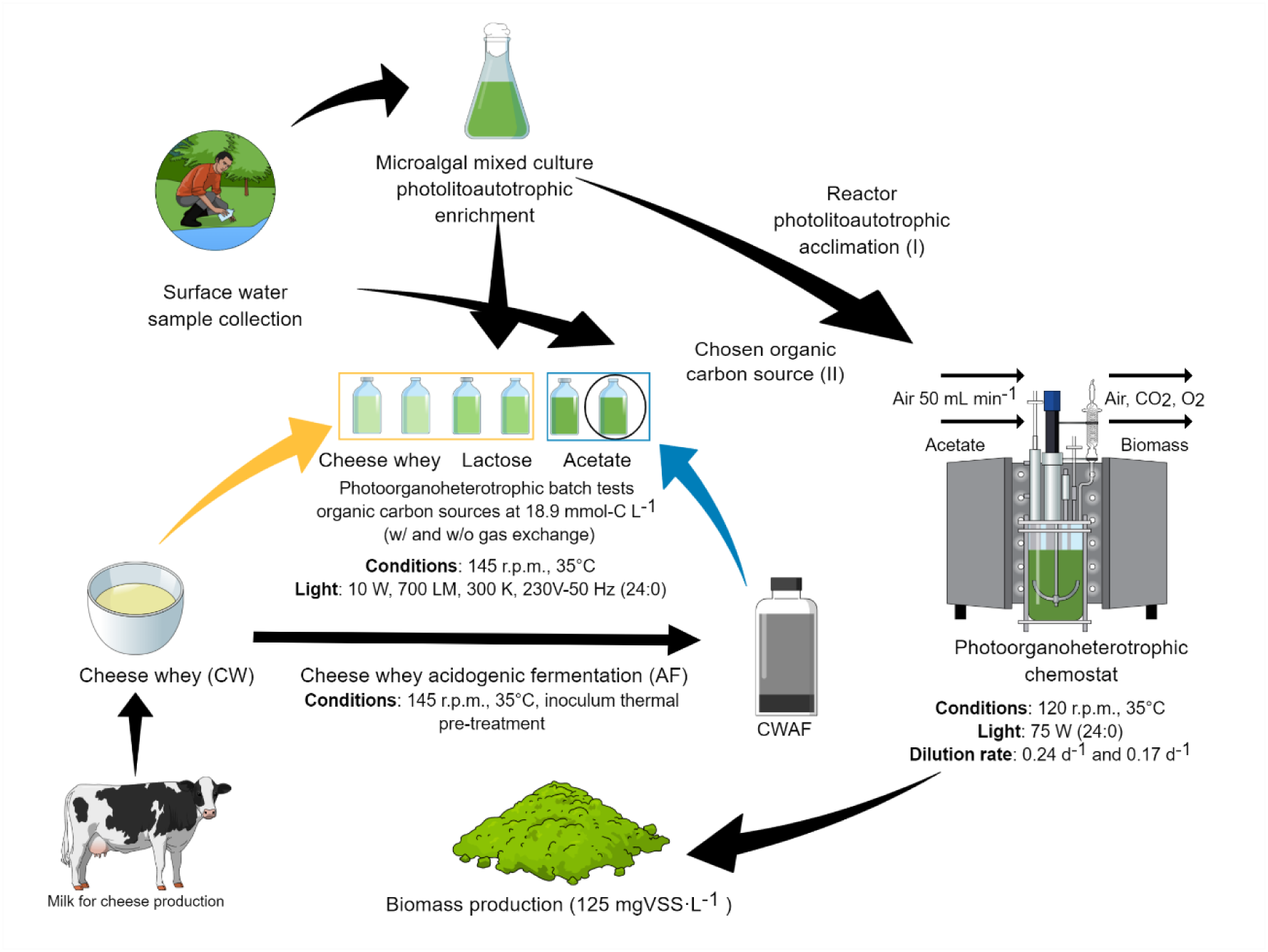

## 1 Introduction

Microalgae are a common and cost-effective polishing step removing nutrients, xenobiotics, and heavy metals from wastewaters foregoing the use of sterile media (Atiku et al., 2016; Beltrán-Rocha et al., 2017; Bohutskyi et al., 2015; Chew et al., 2018; Chiu et al., 2015; de Melo et al., 2018; Fernández-Linares et al., 2017; Gonçalves, Pires, & Simões, 2017; Guldhe et al., 2017; Khan et al., 2019; Kotteswari, Murugesan, & R, 2012; Maizatul et al., 2017; Moreno García et al., 2020; Pahazri et al., 2016; Posadas et al., 2014; Renuka et al., 2015; Singh, Tiwari, & Das, 2016; Solmaz & Mustafa, 2019; Ting et al., 2017; Wang et al., 2016), thriving under stress conditions whilst maintaining their productivity (Ansari et al., 2020). Their photosynthetic activity enables them to sequestrate carbon and decreases wastewater organic and inorganic loads, resulting in biomass that can be either used as a biofertilizer or further processed into biofuels and other products. Consequently, eutrophication can be prevented (Amit & Ghosh, 2018; Bohutskyi et al., 2015; Chiu et al., 2015; Gani et al., 2015; Guldhe et al., 2017; Khan et al., 2019; Kotteswari et al., 2012; Rawat et al., 2011; Renuka et al., 2015; Singh et al., 2016) making phycoremediation a very promising technique for mitigating pollution and environmental impacts (Akthar et al., 2017; Atiku et al., 2016; Beltrán-Rocha et al., 2017; Bohutskyi et al., 2015; Chew et al., 2018; Chiu et al., 2015; de Melo et al., 2018; Fernández-Linares et al., 2017; Gonçalves et al., 2017; Guldhe et al., 2017; Kotteswari et al., 2012; Maizatul et al., 2017; Moreno García et al., 2020; Pahazri et al., 2016; Posadas et al., 2014; Renuka et al., 2015; Singh et al., 2016; Solmaz & Mustafa, 2019; Ting et al., 2017; Wang et al., 2016).

Several studies focus on the use of wastewater for microalgal biomass production (Atiku et al., 2016; Bohutskyi et al., 2015; Cheng et al., 2019; Chiu et al., 2015; Fernández-Linares et al., 2017; Hena, Fatimah, & Tabassum, 2015; Melo et al., 2018; Olguín, 2012; Pahazri et al., 2016; Posadas et al., 2014, 2016; Sahu et al., 2013). Microalgae can be cultivated on arid land (Aratboni et al., 2019; Rösch, Roßmann, & Weickert, 2018; Zeng et al., 2011), brackish or high strength waters (Aslan & Kapdan, 2006; Ho et al., 2011; Jeong, 2003; Kamyab et al., 2015; Ren et al., 2007; Wang et al., 2008; Yi, Han, & Zhuo, 2013; Zhao & Su, 2014), have high photosynthetic efficiency, sequestrate carbon dioxide (Bassi, Saxena, & Aguirre, 2014; Ho et al., 2011; Jeong, 2003; Siregar et al., 2015; Wang et al., 2008; Yi et al., 2013; Zhao & Su, 2014), have different carbon metabolisms (Bassi et al., 2014; Chojnacka & Noworyta, 2004; Crane & Grover, 2010; Godrijan, Drapeau, & Balch, 2020; Subashchandrabose et al., 2013) and an outlet of high added value bioproducts that can be processed into (e.g., biofuels, pigments, omega-3, antioxidants, pigments, lipids, biomass, polyunsaturated fatty acids (PUFAs), polysaccharides, immune modulators, polymers) which can be used by food, aquaculture, nutraceutical, pharmaceutical, cosmetic and energetic industries (Bohutskyi et al., 2015; Chen et al., 2019; Chew et al., 2017; Chiu et al., 2015; D’Imporzano et al., 2018; Fei et al., 2015; Fernández-Linares et al., 2017; Guldhe et al., 2017; Hu et al., 2018; Moreno García et al., 2020; Odjadjare, Mutanda, & Olaniran, 2017; Posten & Schaub, 2009; Pulz & Gross, 2004; Singh et al., 2016).

Microalgae have an outlet of bioproducts of high aggregated value that depending on the substrate used during phycoremediation, there is the possibility of biomass contamination with heavy metals and xenobiotics (Markou et al., 2018). Ergo, coupled anaerobic processes for microalgal mixotrophic biomass growth avoid this contamination by feeding microalgae with only the products of the fermentation of anaerobic digestion, volatile fatty acids.

Anaerobic digestion (AD) is a sustainable option to treat and reuse organic residues, reducing their organic load and concomitantly, converting them into VFAs during fermentation. Anaerobic processes have low energy consumption and the potential to decrease greenhouse gas (GHG) emissions (Boyle, 2012). According to Atasoy et al.(2018), VFAs can serve as raw material to produce some polyhydroxyalkanoates (PHAs), biogas, electricity, and an organic carbon source for the biological removal of phosphorus and nitrogen from wastewater. Microalgae-coupled processes can use VFAs as organic carbon sources derived from anaerobic digestion processes. In our case, this organic carbon source originates from cheese whey acidogenic fermentation processes.

Cheese whey is a product from cheese processing (Archer, 1998; Tsakali et al., 2010) with high organic content (i.e., 50-102 g O_2_ L^−1^ in terms of COD) (Prazeres, Carvalho, & Rivas, 2012; Tsolcha et al., 2016) making cheese whey one of the biggest agricultural pollutants (Kosseva & Webb, 2013; Prazeres et al., 2012; Stamatelatou et al., 2011; Tsolcha et al., 2016). In the year 2022, the forecast for cheese whey production among the world’s biggest producers is 160.67 Mm^3^ y^−1^ although only 58.6% is absorbed by industries (Tsakali et al., 2010; USDA, 2019). The remaining 66.5 Mm^3^ y^−1^ of cheese whey is currently used as animal feed, fertilizers, or in some locations (i.e., developing and developing countries and remote locations), it can be illegally discharged into water bodies (Tsakali et al., 2010).

The acidogenic fermentation of cheese whey produces mainly acetate (Hassan & Nelson, 2012). Phototrophic organisms can convert electromagnetic energy into chemical energy to maintain cellular growth (Overmann & Garcia-Pichel, 2013). Microalgae (i.e., microalgae and cyanobacteria) are phototrophic organisms capable of performing mixotrophy (Mondal et al., 2016; Salati et al., 2017).

Some microalgae can display heterotrophic and mixotrophic metabolisms (Bouarab, Dauta, & Loudiki, 2004; Hu et al., 2018; Mondal et al., 2016). The mixotrophic regime is thought to be the sum of both photolithoautotrophic and chemoorganoheterotrophic regimes (Bouarab et al., 2004). In mixotrophic microalgae mixed cultures coupled with cheese whey anaerobic digestion processes, the organic carbon source can come from the lactose, proteins, and fats present in cheese whey or, as we propose it can come as acetate, the main VFA produced in cheese whey anaerobic processes.

Another way to obtain organic carbon sources is through symbiotic relationships within ordinary heterotrophic organisms (OHO). They will degrade organic matter into carbon dioxide and in return, microalgae will produce oxygen in photosynthesis, which is essential for aerobic bacterial life (Yao et al., 2018). However, competition for substrate and predatism can be an issue in mixotrophic cultures, decreasing microalgae organic carbon uptake (Chen, Zhao, & Qi, 2015; Ramanan et al., 2016). Hence, it can influence biomass production. Mixotrophic cultures also have an economic advantage over axenic cultures as they do not require sterilization of substrate and equipment as do pure cultures to avoid contamination (Chen et al., 2015).

Our work aims to understand the ecological engineering of green photoorganoheterotrophs and their mechanisms of organic carbon uptake. We also target the best reactor regimes and parameters (i.e., dilution and hydraulic retention time) for green phototroph selection. In addition, we attempt to comprehend the symbiotic relationships between different guilds of microorganisms for upscaling the process.

## 2 Materials and Methods

### 2.1 Enrichment of mother culture

Surface water was sampled from a channel 51°59’23.3”N 4°22’53.8”E (Ruiven, 2628 CN Delft, NL). A mother microalgal culture was enriched from this sample using Alleńs Blue-Green algal medium, BG-11 (Allen, 1968; Allen & Stanier, 1968; Ripka et al., 1979), in a 1:15 ratio. Acclimation lasted 15 days. Every two weeks, the mother culture was reinoculated.

The culture was initially grown photolitoautrotophicaly in an incubator (shaker KS-15, hood TS-15, Edmund Bühler GmbH, Germany) at 145 rpm mixing and mesophilic temperature of 35°C. A led lamp (10 W, 700 LM, 300K, 230V-50 Hz, GAMMA, The Netherlands) provided continuous illumination of 80 μmol photons m^−2^ s^−1^. Culture flasks used a sterile cotton stopper allowing gas exchange between media and atmosphere. Once culture was established, it served as an inoculum for all experiments.

### 2.2 Study of different phototropism regimes in batches

**Table 1** describes the experimental set-up for photolitoautotrophic biomass growth with different nitrogen sources and gas exchange conditions and photoorganoheterotrophic microorganisms’ selection under different carbon source sections.

**Table 1.**
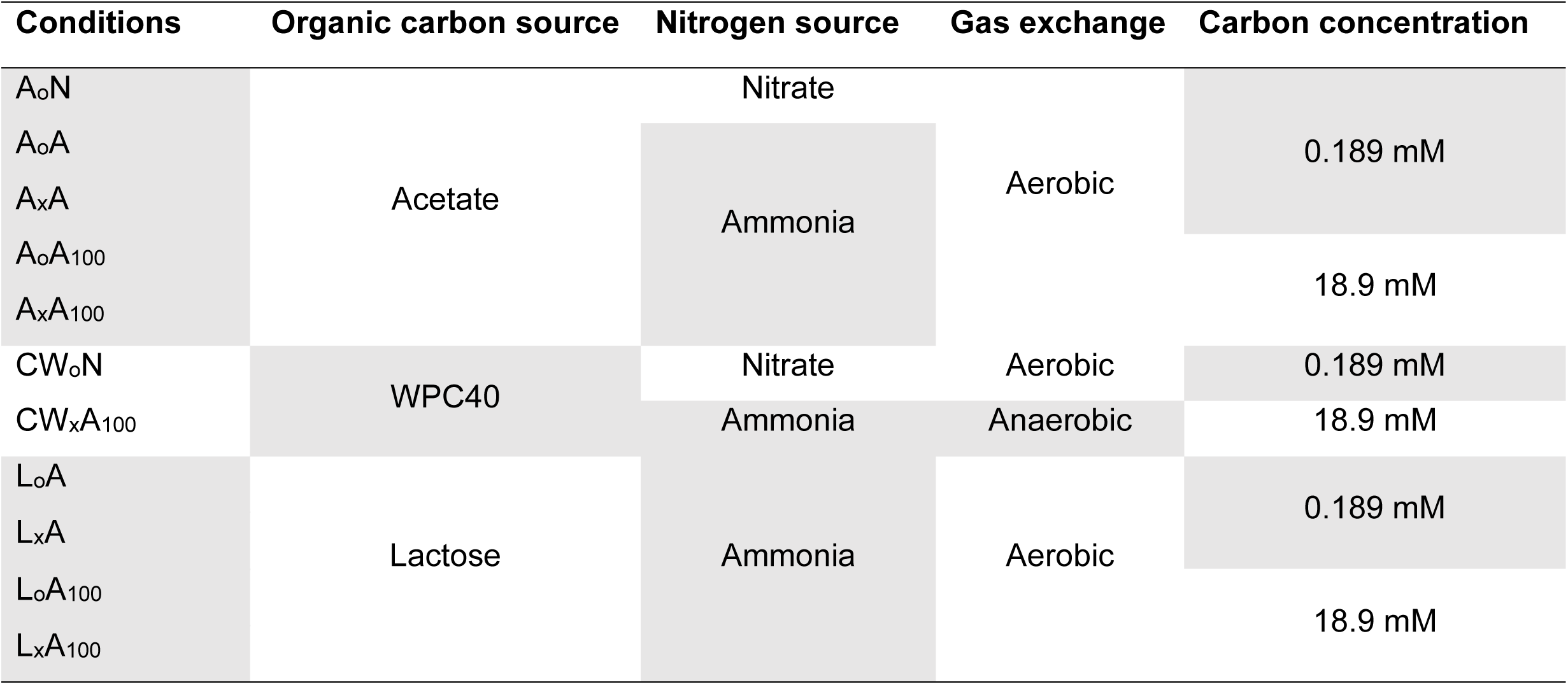
Set up experiments for photolitoautotrophic biomass growth with different nitrogen sources and gas exchange conditions and photoorganoheterotrophic microorganisms’ selection under different carbon source sections. Table 1 summarizes the experiments regarding their organic carbon and nitrogen source, aerobic or anaerobic conditions, and carbon concentration. An initial organic carbon concentration of 0.189 mM was equivalent to the inorganic carbon concentration in the BG-11 medium whereas, conditions with a concentration of 18.9 mM C L^−1^ were 100-fold BG-11 conditions. The analysis encompassed solid series, cell counting and viability, pH measurements, chlorophyll extraction, nutrient consumption, and microscopy.

#### 2.2.1 Photolitoautotrophic growth with different nitrogen sources and gas exchange conditions

Cultivation was performed in borosilicate vials (100 mL) using a modified BG-11 medium heated at 90°C for whole carbon source (CO_3_^2-^) dissolution. The nitrate (NaNO_3_) content of the medium was substituted by ammonium (NH_4_Cl) in equal mols (17.6 mmol N L^−1^). Gas exchange was controlled by using open and sealed bottles which were sealed with rubber stoppers and aluminum crimp caps.

Vials were sparged with nitrogen to ensure initial anaerobic conditions. Open bottles had a sterile cotton stopper to allow gas exchange. Measurements consisted of biomass concentration as volatile suspended solids (VSS), nutrient (N and P) consumption, pH variation, and microscope visualization. The experiment lasted for 30 days.

#### 2.2.2 Selection of photoorganoheterotrophic microorganisms by different carbon sources

A photoorganoheterotrophic regime was applied to the mother culture using different organic carbon sources such as acetate, lactose, and 40% demineralized whey powder (WPC40). BG-11 medium was used as a substrate where each carbon source was substituted in the medium by 0.189 mM C L^−1^ and then by 100-fold higher as the original recipe. When the carbon source was substituted by WPC40, a total organic carbon (TOC) assay was performed to measure its carbon content. Cultivation was performed in 250 mL Erlenmeyer in a 1:15 ratio of inoculum and medium. Nitrate (NaNO_3_) content in the medium was substituted by ammonium (NH_4_Cl) in equal mols (17.6 mmol-N L^−1^) as in the medium.

Sealed vials were sparged with nitrogen to ensure initial anaerobic conditions. Open bottles had a sterile cotton stopper to allow gas exchange. The experiment lasted for 10 days. An additional experiment to verify the effect of organic carbon source in biomass and pigment concentration was performed in microplates using WPC40 and acetate. Different concentrations of organic carbon (0.221, 0.947, 2.524, 6.31, 12.621 to 25.241 mM) in the forms of acetate and WPC40 replaced the carbon concentration in BG-11 medium resulting in C:N:P ratios of 1:101:1, 5:101:1, 14:101:1, 36:101:1, 72:101:1, 144:101:1. Absorbance was measured over the visible light spectrum using a microplate reader (Synergy™ HTX, BioTek, USA). The experiment lasted 10 days.

#### 2.2.3 Photoorganoheterotrophic growth of microalgae in a continuous-flow photobioreactor

An initial photolitoautotrophic regime on BG-11 medium was imposed on microalgae until growth stabilization in batch mode with a 0.24 ^−1^ dilution rate. This experiment lasted 12 days. Due to incomplete carbon dissolution, BG-11 was then heated to 90°C leading to complete carbon dissolution. Biomass growth and nutrient consumption this time was monitored for 7 days.

The reactor regime changed to photoorganoheterotrophic conditions using 18.9 mmol C L^−1^ of acetate in continuous-flow mode as established in section 2.2.2. The experiment was scaled up in a 3.5 L continuous-flow stirred-tank photobioreactor (CSTR-PBR) with a 2 L working volume. An initial photolitoautotrophic regime on BG-11 medium was imposed on microalgae until growth stabilization in batch mode. Additional nutrients were provided according to BG-11 medium, except for the nitrogen source that was replaced by NH_4_Cl in equal mols (17.6 mmol-N L^−1^) as stated in the medium.

Both reactor regimes were continuously irradiated with a 75 W white light jacket (400-700 nm, PhotoBioSim, Designinnova, India) at 96 μmol photons m^−2^ s^−1^ and run at a mesophilic temperature of 35°C. Photobioreactor in batch mode started with a dilution rate of 0.24 d^−1^. Hence, the initial dilution rate of the chemostat was also 0.24 d^−1^ changing to 0.17 d^−1^ after a few days. Argon was sparged at 50 mL min^−1^. Gas flow and off-gas CO_2_ and O_2_ measurements were monitored. Polyisocyanurate (PIR) boards were used to serve as an isolation hood so no external light interfered with CSTR-PBR light conditions. An anchor paddle provided CSTR_PBR mixing at 120 r.p.m. and prevented biofilm formation. Biomass growth was monitored daily until a steady state was reached with nutrient consumption afterward. **Figure 1** shows the reactor setup and the isolation hood.

**Figure 1.**
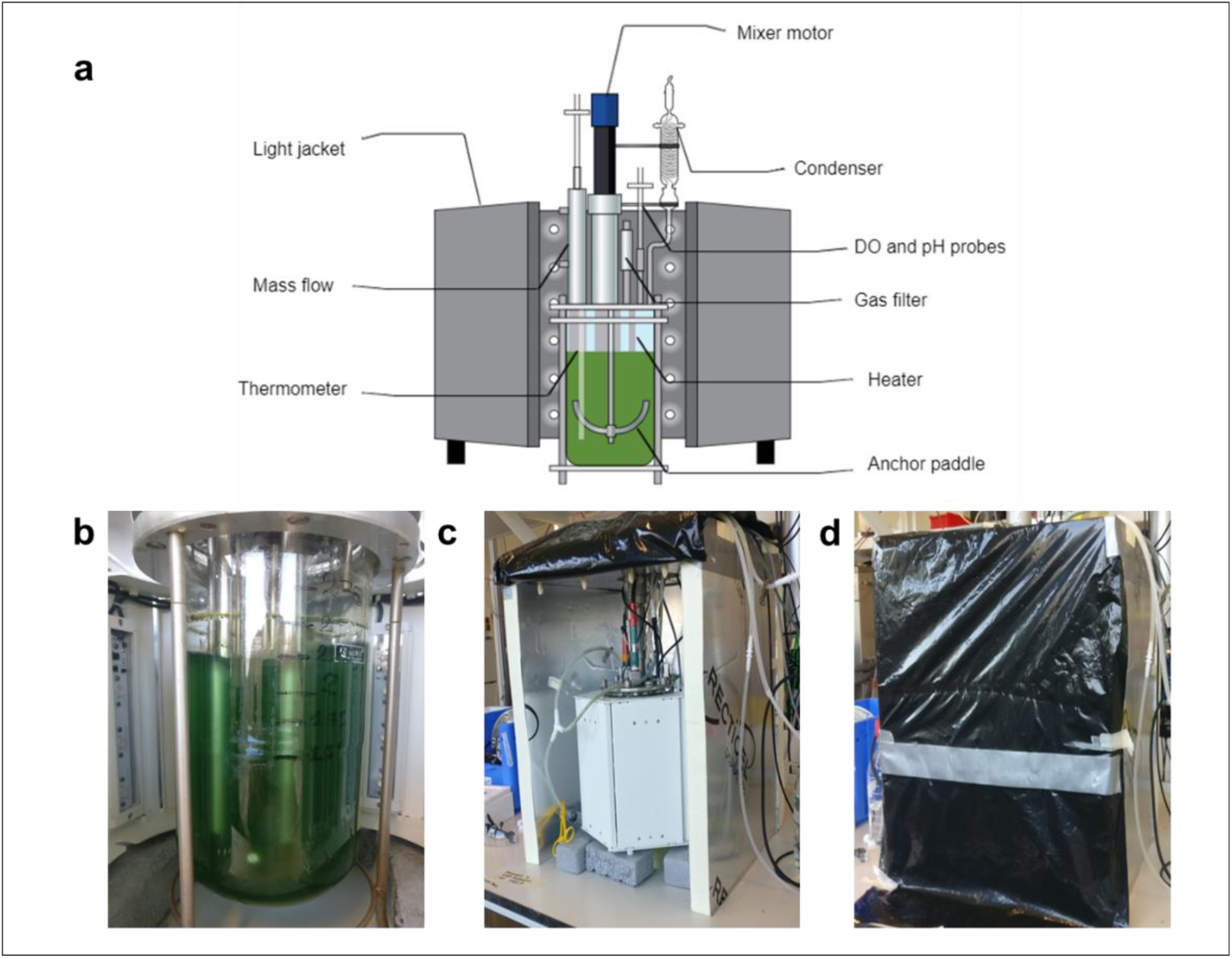
**a**. Photobioreactor set up with a light jacket surrounding it. The temperature was kept at 35°C and argon was injected in the reactor. The incoming gas was controlled by a mass flow controller, while the off-gas was cooled by a condenser and filtered with a cotton filter before it was sent to the off-gas analyzer. DO, temperature, and pH were continuously measured in all set-ups. For the chemostat regime, influent and effluent flows were regulated by pumps. Acid and base were dosed to maintain the pH at 7. **b.** Picture of the reactor the a and light jacket. **c**. Picture of the PIR-boards hood built to avoid the influences of external light and temperature. **d.** PIR-boards hood had a black plastic cover that enabled access and hinder the entry of external light.

### 2.3 Physical and chemical analysis

#### 2.3.1 General analysis

The following analysis was common to most experiments, or else specified. Ammonium, nitrite, nitrate, and phosphate content were either measured by manual spectrometric method (ISO 7150-1:1984) (ISO, 1984) or by discrete analysis systems with photometric detection (ISO 15923-1:2013) (ISO, 2013).

Total organic carbon (TOC) was measured by combustion catalytic oxidation method at 720°C – standard methods 5310B (APHA, 2005) (APHA, 2005), and total nitrogen (TN) was also analyzed at 720°C by catalytic thermal decomposition method/chemiluminescence – Standard Method 5310B(APHA, 2005). Both analyses were performed on the first and last day of the experiments.

In most experiments, dissolved oxygen (DO) and pH were measured daily by electrometry and potentiometry, respectively (APHA,2005) (APHA, 2005). Gas flow was regulated by a mass flow controller (0154 Microprocessor Flow Control & Read Out Unit, Brooks Instruments, USA). Argon was used as the off-gas carrier at 50 mL min ^−1^ and CO_2_ and O_2_ off-gas measurements were performed by an online gas analyzer (NGA 2000, Rosemount, USA).

Solids series were analyzed according to Standard Method 2540 (APHA, 2005)(APHA, 2005). For mixed cultures observation, about 1 µL of unstained samples were fixed with alcohol 70% (v/v) to stop motility and observed by phase contrast microscopy using Axioplan 2 (Zeiss, Germany).

#### 2.3.2 Acetate and carbohydrates consumption

VFA (acetate) and sugar (lactose, glucose, and galactose) consumption were monitored daily. A sample containing a volume of 2 mL was filtered with a 0.45 µm pore size disposable sterile polytetrafluoroethylene polymer (PTFE) syringe filter with needles for filtration. About 1 mL of sample was then detected by high-performance liquid chromatography (HPLC - Waters, USA), with a BioRad Aminex separation column (Biorad, USA HPX-87H) and a BioRad Cation-H refill cartridge guard column. Chromatographer was equipped with a 717 plus Autosampler, a 2489 UV/Vis Detector, and a 2414 Refractive Index Detector (Waters Corporation, USA). The mobile phase was a solution of 1.5 mM phosphoric acid (H_3_PO_4_) at 60°C and a flow rate of 0.6 mL min^−1^. Glucose consumption was also monitored according to the phenol-sulphuric method described by Dubois (1956).

#### 2.3.3 Cell viability and biomass analysis

Cell viability was monitored by methylene blue (MB) as proposed by Bicas et al.(2015). Two samples of 3-5 mL were taken from the culture, where one of them was heated to 90°C in a water bath for 5 minutes. After the heated sample reached room temperature, both samples were centrifuged for 3 minutes at 4000 g min^−1^.

The supernatant was discarded and pellet resuspended into 4 mL of methylene blue (MB) solution (85 μM of MB in phosphate buffer solution 0.1 M, pH 7.4), followed vortexing and centrifugation at 4000 g min^−1^. Supernatant was read at 664 nm and compared to the absorbance of MB solution at the same wavelength. The uptake of MB was measured with **equation 1**:

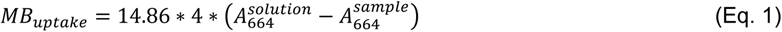

Cell viability was given by considering the ratio between the MB uptake of both raw and boiled samples.

Biomass concentration was determined by optical density at 750 nm (A_750_) and the constant term of calibration (q) using a spectrophotometer (DR3900, Hach Lange, USA). A calibration curve was created for conversion of absorbance to grams of biomass per liter using **equation 2**:

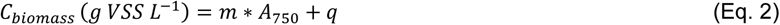

Other conversions such as maximum growth rates, yields, production rates, and biomass-specific rates of substrate were calculated according to the following **equations 3** to **6**, respectively:

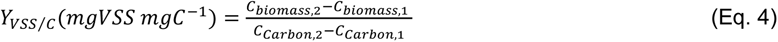

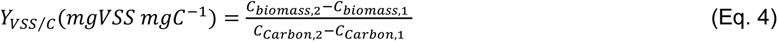

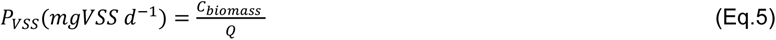

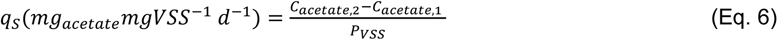

#### 2.3.4 Pigments extraction and quantification

An adaptation of Shoaf and Lium (1976) was used for pigment extraction and quantification. Samples of 2.5 mL were collected, and centrifuged at 13,300 g min^−1^ for 3 minutes. Supernatant was discarded and the pellet was resuspended in a 1:1 (v/v) solution of dimethyl sulfoxide (DMSO) and 80% acetone (80%). Eppendorffs containing samples were closed with parafilm to avoid evaporation and covered in foil paper to avoid any light interference. Samples were placed at 4°C for 24 hours. The following day, samples were centrifuged (13,300 g min^−1^, 3 minutes) and the absorbance spectrum (350 to 750 nm, DR3900, Hach Lange, USA) was measured. A DMSO:acetone solution was used as blank. Measurements were performed in a quartz cuvette to avoid any fog common on polystyrene disposable cuvettes.

Chlorophylls a, b, and carotenoid contents were calculated according to Wellburn(Wellburn, 1994). Specific sets of equations were used to calculate contends in 80% acetone solution (**equations 7, 8**, and **9**) and DMSO (**equations 10, 11**, and **12**). After calculation, the result was given as an average of results from both solvents. According to Arnon et al. (1949) and Speziale et al. (1984), averaging results is plausible given the negligible difference between equations.

##### Acetone

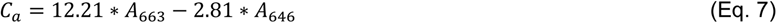

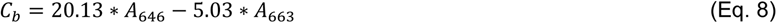

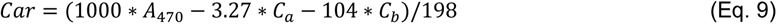

##### DMSO

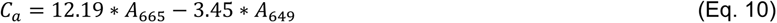

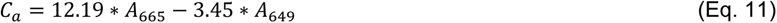

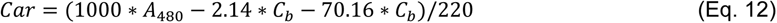

In addition to pigment extraction, a wavelength scan over the visible light spectrum (330 to 900 nm) was performed with a spectrophotometer (DR3900, Hach, Germany) to monitor the evolution of pigment and biomass content over the experiment.

### 2.4 Microbial composition gene amplicon sequencing analysis

DNA extractions for 16S and 18S rRNA were performed to identify prokaryotes and eukaryotes microorganisms in different phototrophic regimes (photolitoautotrophic, photolitoheterotrophic, and photoorganoheterotrophic) in batches and CSTR-PBR with DNeasy® UltraClean® Microbial Kit (QIAGEN, Germany).

Both DNA extractions’ concentration and quality were measured by fluorescence using an Invitrogen Qubit 4.0 (Thermo Fisher Scientific, USA) and 200 μL samples of concentrated DNA (ranging from 13.9 to 95 ng μL^−1^) were stored at −20°C. Novogene Co., Ltd. (China) was responsible for the amplicon sequencing. At Novogene, DNA concentration and purity were evaluated on 1% agarose gels, and depending on the concentration, samples can be diluted to 1 ng μL^−1^ with mili-Q water.

Selected regions for 16S rRNA (V3-V4) and 18S rRNA (V4) were amplified by polymerase chain reaction (PCR) for library preparation. For 16S rRNA, the regions V3-V4 had the primer forward 341F (5’-CCTAYGGGRBGCASCAG-3’) and reverse 806R (5’-GGACTACNNGGGTATCTAAT-3’) according to Takahashi et al.(Takahashi et al., 2014). In 18S rRNA amplification, region V4 had the primer forward 528F (5’-GCGGTAATTCCAGCTCCAA-3’) and reverse 706R (5’-AATCCRAGAATTTCACCTCT-3’) according to Cheung et al.(Cheung et al., 2010) and Lutz et al. (Lutz et al., 2015)

Sequencing libraries were created using NEB Next Ultra ^TM^ DNA Library Prep Kit for Illumina^®^ (QIAGEN, Germany). Library quality was evaluated with a Qubit 2.0 Fluorometer (Thermo Scientific, USA) and Agilent Bioanalyzer 2100 system (Agilent, USA). The library was then sequenced on an Illumina platform (Illumina, USA) and 250 bp paired-end reads were generated.

PCR reactions were performed by 30μL reactions using 15μL of Phusion® High-Fidelity PCR Master Mix (New England Biolabs, USA), 0.2 μM of primers

Set, and about 10 ng of template DNA. Thermal cycling had an initial denaturation (98°C, 1 minute), followed by 30 cycles of denaturation (98℃, 10 seconds), annealing (50℃, 30 seconds), elongation (72℃, 1 minute,) and a 5 minutes step at 72℃. PCR products were quantified and qualified, then mixed in equidensity ratios. The mixed PCR products were purified with a GeneJET Gel Extraction Kit (Thermo Scientific, USA).

Data analysis constituted of identification of nucleotides (base calling), removal of chimeric sequences, and generation of operational taxonomic units (OTU) with ≥ 97% identity. OTUs were also analyzed using the SINA (v.1.2.11) aligner based on the global SILVA alignment for rRNA genes.

## 3 Results and Discussion

### 3.1 Photolithoautotrophs are set in their ways

#### 3.1.1 Biomass growth and cell viability: thriving in BG-11

Photolithoautotrophic culture on BG-11 and gas exchange allowance displayed an increase in three units of initial pH of 7.0. Usually, carbon dioxide dissolution decreases medium pH and carbonic acid formation. Consequently, bicarbonate is formed with hydrogen ions dissolution and subsequent carbonate ion formation. However, the increase observed in pH can result from hydrogen ions absorption during microalgae photosynthesis and denitrification given the non-axenic mother culture. Increased medium turbidity and reduction in nitrogen and phosphorus levels indicated microorganisms’ absorption for anabolic processes.

Biomass concentration was verified by dry weight measurement, and cell viability. Despite the pH increase, all measurements indicated biomass growth. Dry weight as volatile suspended solids (VSS) were of about 650 mg VSS L^−1^, indicating an increase of about 13-fold the initial biomass concentration. Methylene blue (MB) cell viability measurements showed a decrease in viable cells after the tenth day reaching 17% by the end of measurements. Decrease in absorbance is proportional to decrease in cell viability. **Figure 2** shows biomass concentration increase and cell viability.

**Figure 2.**
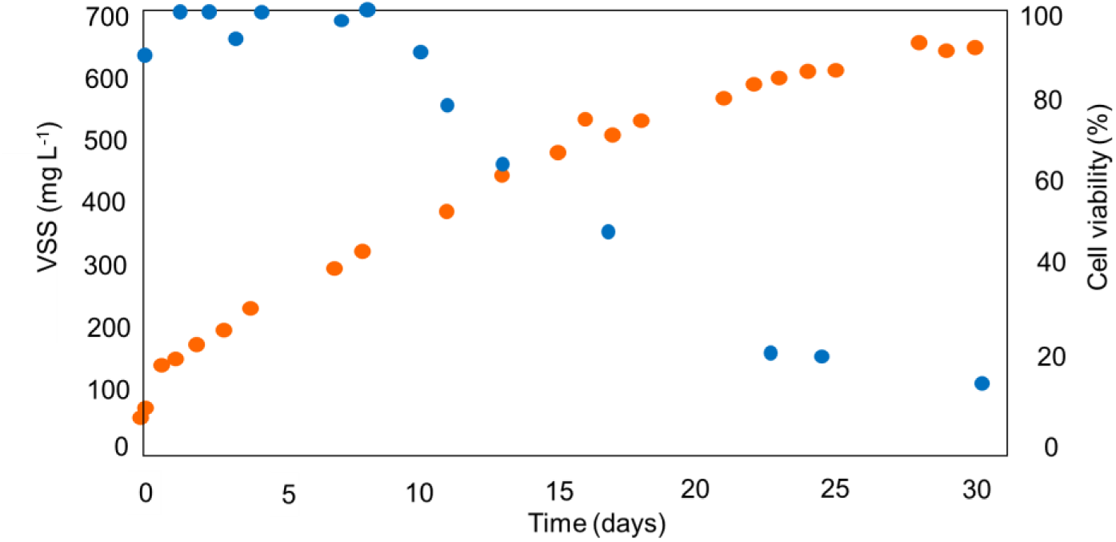
Photolitoautotrophic culture biomass concentration (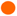) and cell viability (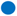).

#### 3.1.2 Pigment content increase is consistent with biomass growth

Pigments were identified in the following culture absorbance spectra peaks: 440, 630 and 680 nm which corresponded to chlorophyll *a* and *b*. Sample absorbance spectrum are shown in **figure 3**. The solvents used for pigment extraction, DMSO and acetone. The blank used was distilled water. Pigments absorbance peaks might have been slightly lowered and shifted to the left due to the difference on composition of medium and blank.

**Figure 3.**
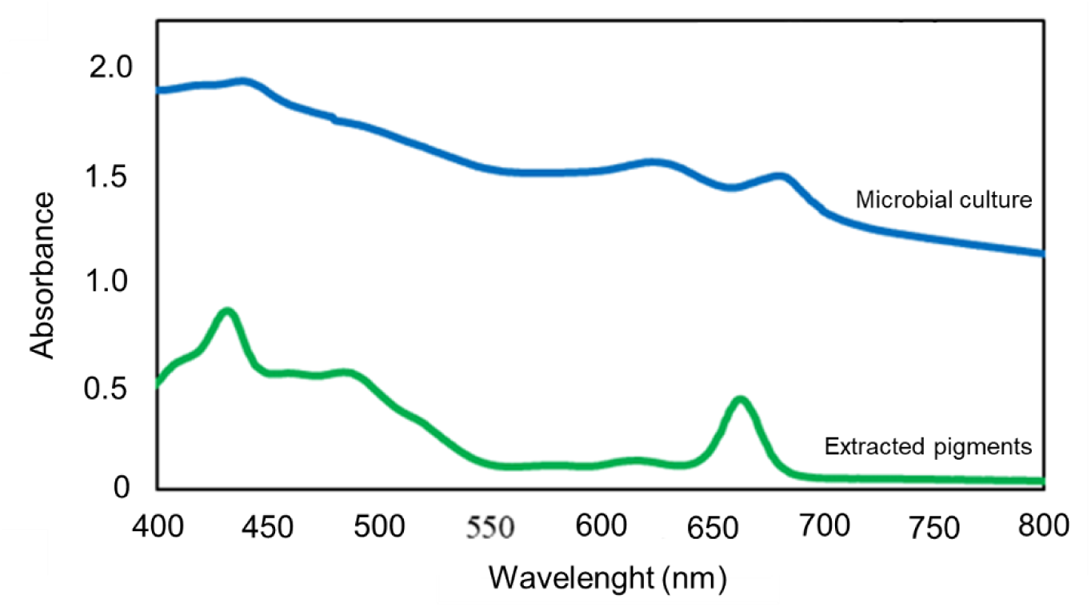
Absorbance spectra of the microbial culture and pigments extracted with DMSO/Acetone measured after 13^th^ day of cultivation. The bottom curve in green regards the extracted pigments, whereas the upper curve refers to the microbial culture.

Pigments content was directly proportional to biomass concentration and increasing linearly until the last measurement after the exponential growth phase on the 16^th^ day. Biomass specific pigment content also increased, as shown in **table 2**.

**Table 2.**
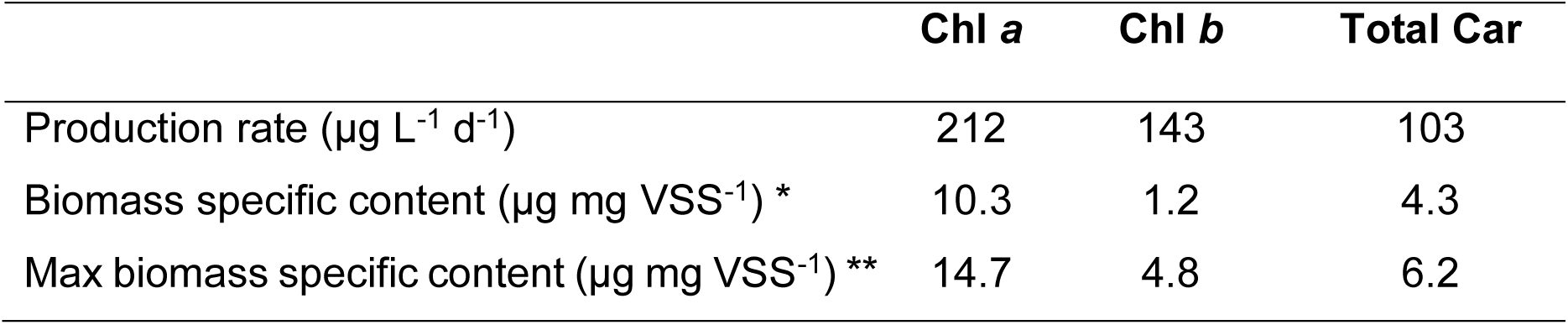
Pigments production according to their production rate, biomass specific content and the maximum biomass specific content. *Their production rate was constant after the 5^th^ day of measurement. ** The maximum biomass specific content refers to the end of the experiment. *Chl a* stands for chlorophyll a, *Chl b* for chlorophyll b and *Car* for carotenoids.

|  | Chl a | Chl b | Total Car |
| --- | --- | --- | --- |
| Production rate ($\mu\text{g L}^{-1} \text{d}^{-1}$) | 212 | 143 | 103 |
| Biomass specific content ($\mu\text{g mg VSS}^{-1}$) * | 10.3 | 1.2 | 4.3 |
| Max biomass specific content ($\mu\text{g mg VSS}^{-1}$) ** | 14.7 | 4.8 | 6.2 |

#### 3.1.3 Changes in carbon and nitrogen sources are not beneficial to photoaautolitotrophic culture

Carbon and nitrogen changes in mother culture with BG-11 medium were proposed to improve cell growth and biomass concentration. Initially, incomplete medium carbonate dissolution, which was thought to hinder biomass growth, was then addressed by heating the solution of Na_2_CO_3_ before adding it to the medium. However, carbonate dissolution by heat caused a decrease of 48% in cell productivity and 33% in biomass concentration within the first 10 days of experiment.

In addition, a drastic increase in pH overnight from 7.0 to 10.0 indicated that most microorganisms in the photolitoautotrophic community were also intolerant to alkaline pH. The same decrease in cell productivity, biomass concentration happened when replacing the nitrogen source from nitrate (NaNO_3_) to ammonium (NH_4_Cl). However, it could be observed a decrease in pH. The aims behind these changes were to limit the action of denitrifiers and boost acidogenic fermentation, but they also led to lower biomass concentration. The proposed modifications in BG-11 medium did not show positive changes in photolitoautotrophic cultures. **Table 3** shows the alterations in cell growth and biomass concentration consequence of changes in nitrogen and carbon sources.

**Table 3.**
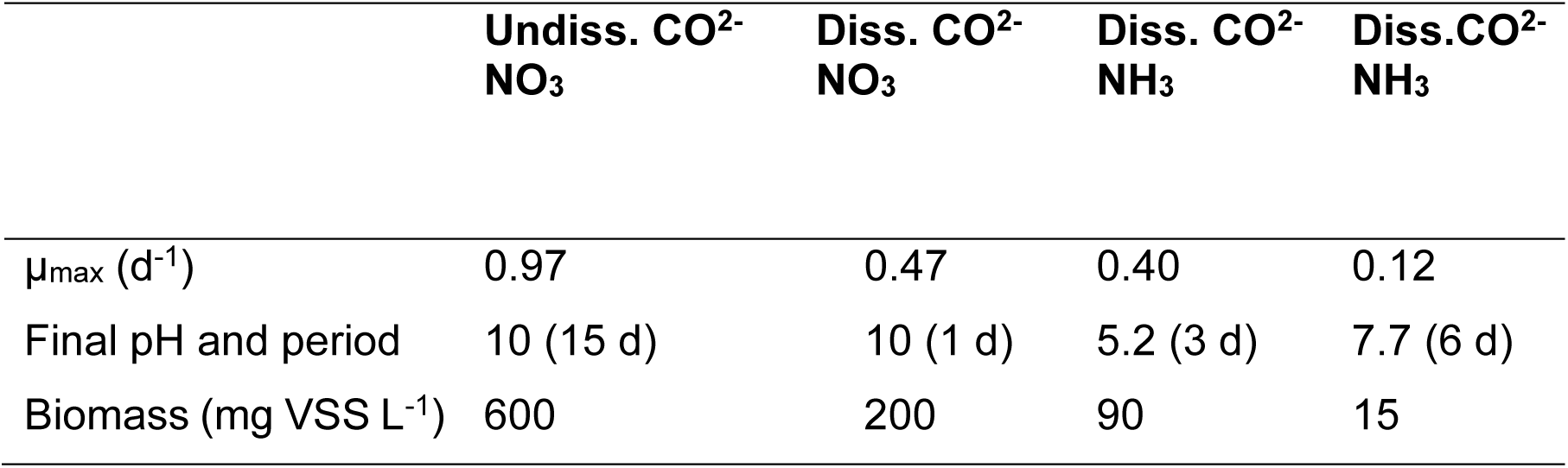
Variations in μ_max_, biomass increase and pH evolution according different media and aerobic conditions. The number of days in which pH reached a new value are in parentheses. Initial BG-11 pH is 7.0. (Undiss) stands for undissolved and (Diss) for dissolved.

|  | Undiss. CO<sub>2</sub><sup>-</sup><br>NO<sub>3</sub> | Diss. CO<sub>2</sub><sup>-</sup><br>NO<sub>3</sub> | Diss. CO<sub>2</sub><sup>-</sup><br>NH<sub>3</sub> | Diss.CO<sub>2</sub><sup>-</sup><br>NH<sub>3</sub> |
| --- | --- | --- | --- | --- |
| $\mu_{\max}$ (d<sup>-1</sup>) | 0.97 | 0.47 | 0.40 | 0.12 |
| Final pH and period | 10 (15 d) | 10 (1 d) | 5.2 (3 d) | 7.7 (6 d) |
| Biomass (mg VSS L<sup>-1</sup>) | 600 | 200 | 90 | 15 |

#### 3.1.4 Visualization and identification of photolitoautotrophic cultures revealed microorganism’s diversity

Observations in phase-contrast microscopy revealed great microorganism diversity with an abundance of green round-shaped cells of 3-4 μm diameter, mostly in aggregates. Other types of microorganisms were not as plentiful. **Figure 4** shows the composition of microorganisms in photolitoautotrophic cultures.

**Figure 4.**
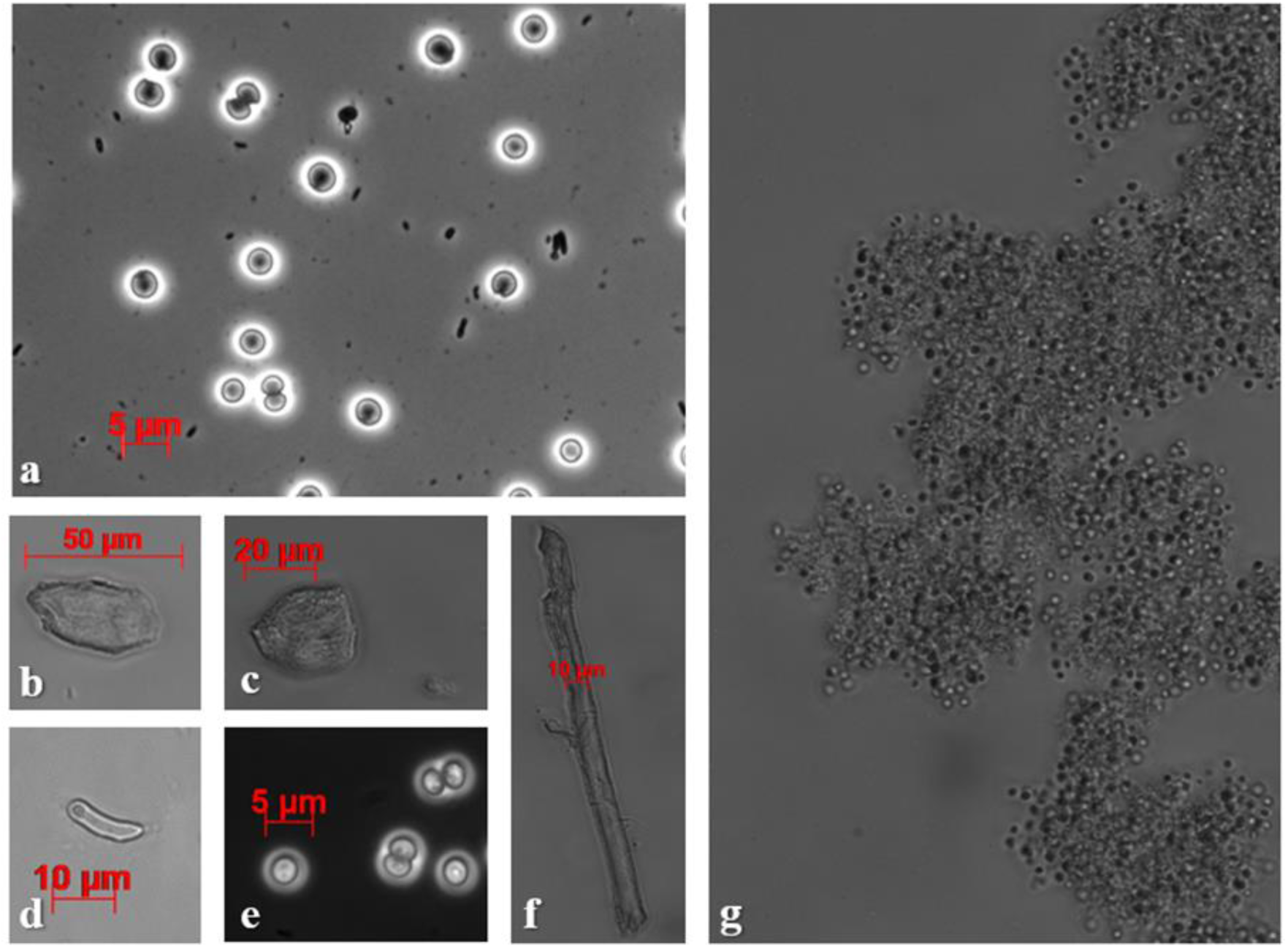
(***a-g***) Photolitoautotrophic culture bacterial community. Images *a*, *e* and *g* depicted the most abundant 3-4 μm diameter round-shaped cells. They tend to form aggregates as shown on image ***g***. Larger microorganisms were found on images ***b*, *c*, *d*** and ***f***, whereas smaller bacteria can be observed on the background of image ***a***.

As for the identification of microbial populations, 16S rRNA gene amplicon sequencing confirmed microorganisms diversity as seen through phase-contrast microscopy. Cyanobacteria, mostly Cyanobium PCC-6307 sp. and Synechocystis PCC-6803 sp., corresponded to 10-20% of prokaryotic community. The phototrophic purple non-sulfur bacteria (PNSB) *Rhodobacter* sp. corresponded to 10% of initial community but they were outcompeted along the cultivation period.

All bacteria presented in initial and final cultivation were gram-negative and most of them were able to oxidize nitrate. Denitrifier bacteria are mostly facultative anaerobic chemoorganoheterotrophic microorganisms. In the BG-11 photolitoautotrophic mother culture, NaNO_3_ was the nitrogen source. This suggests that these bacteria were able to also use the inorganic nitrogen present in the medium and the dissolved oxygen as terminal e-acceptor.

In the 18S rRNA gene amplicon sequencing, in the final sample there was a visible decrease of green microalgae genera (i.e., *Scenedesmus* sp.*, Desmodesmus* sp. And Trebouxioupheceae) found in the initial culture to an increase of unidentified Chlorophyceae. The increase of the genus Vermamoeba indicates lack of competition among microorganisms or even their predatory activities which enabled other genus of bacteria to thrive in throughout culturing. **Figure 5** shows the prokaryotic and eukaryotic diversity between initial and final samples of mother-culture.

**Figure 5.**
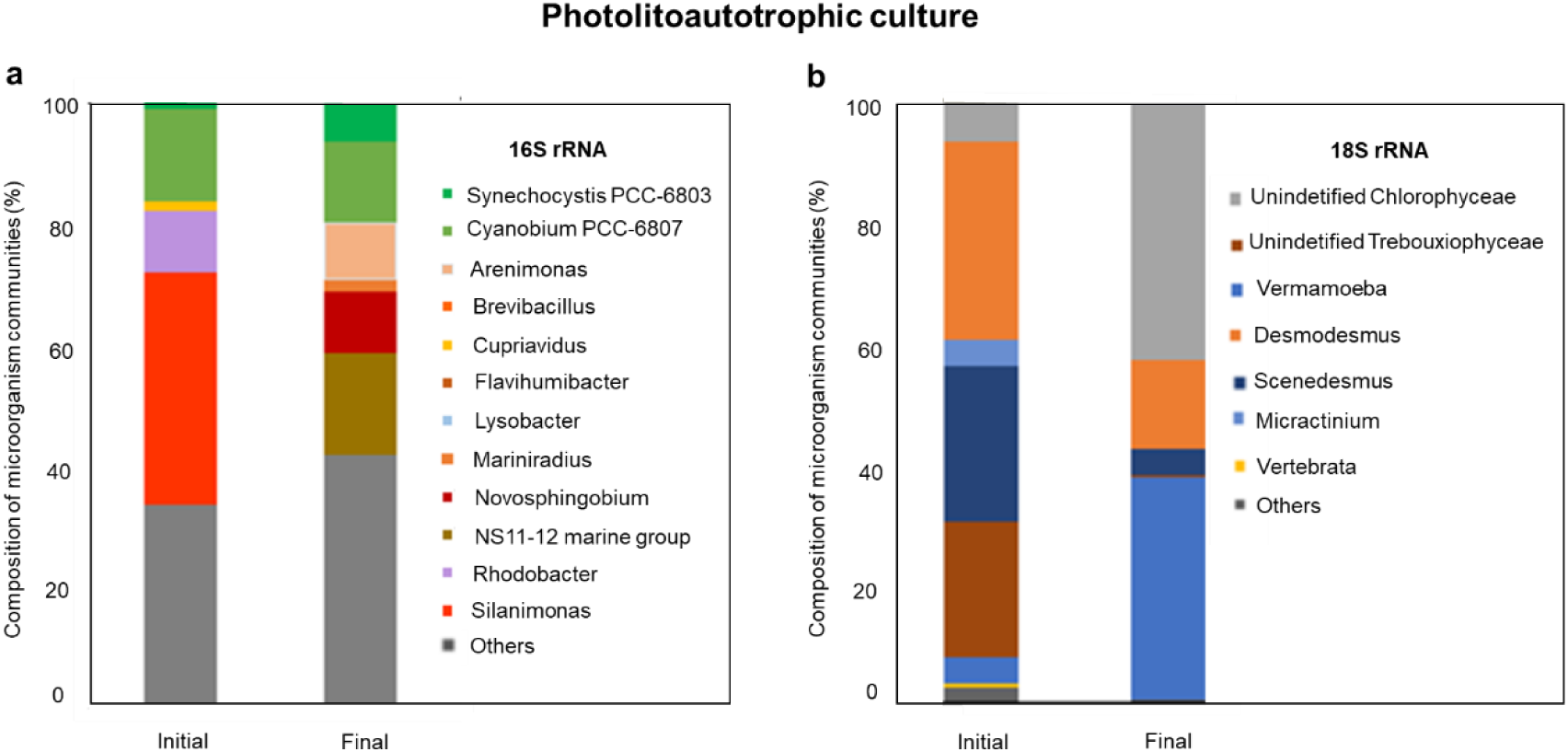
Composition of microbial communities. Figure **A** shows the results from 16S rRNA amplicon gene sequencing region V3-V4 and figure **B**, the results from 18S rRNA amplicon gene sequencing region V4. Only genera comprising more than 2% of the composition of communities are depicted in the graph. The remaining were classified as ‘others’.

### 3.2 Influence of different organic carbon sources in photoorganoheterotrophic microrganisms selection

#### 3.2.1 High concentration of organic substrates enables green phototrophs selection in mixed cultures

The medium chosen for mixed culture enrichment was BG-11 and substitution of inorganic carbon source to organic ones (e.g., WPC40, lactose and acetate) was done with 0.189 mM C L^−1^ and with 100-fold the concentration in the medium, 18.9 mM C L^−1^. Physico-chemical, morphologic and ecologic patterns displayed by 0.189 mM C L^−1^ open bottle cultures grown on acetate and WPC40 as carbon sources, and NaNO_3_ as nitrogen source were similar to the photolitoautotrophic culture.

In both 0.189 mM C L^−1^ close and open bottles batches, exchanging nitrogen source led to variations in cell growth patterns. Open bottles batches with acetate and lactose as organic carbon source had 0.46 d^−1^ and 0.44 d^−1^ growth rates, respectively. These rates were similar to the 0.4 d^−1^ growth rate of photolitoautotrophic conditions. A similar microorganism community pattern to photolitoautotrophic culture can be observed in **figure 6**. An additional pH decreased culture inorganic carbon uptake, since atmospheric CO_2_ absorption was limited with subsequent carbonate saturation. Consequently, it has halted biomass growth.

**Figure 6.**
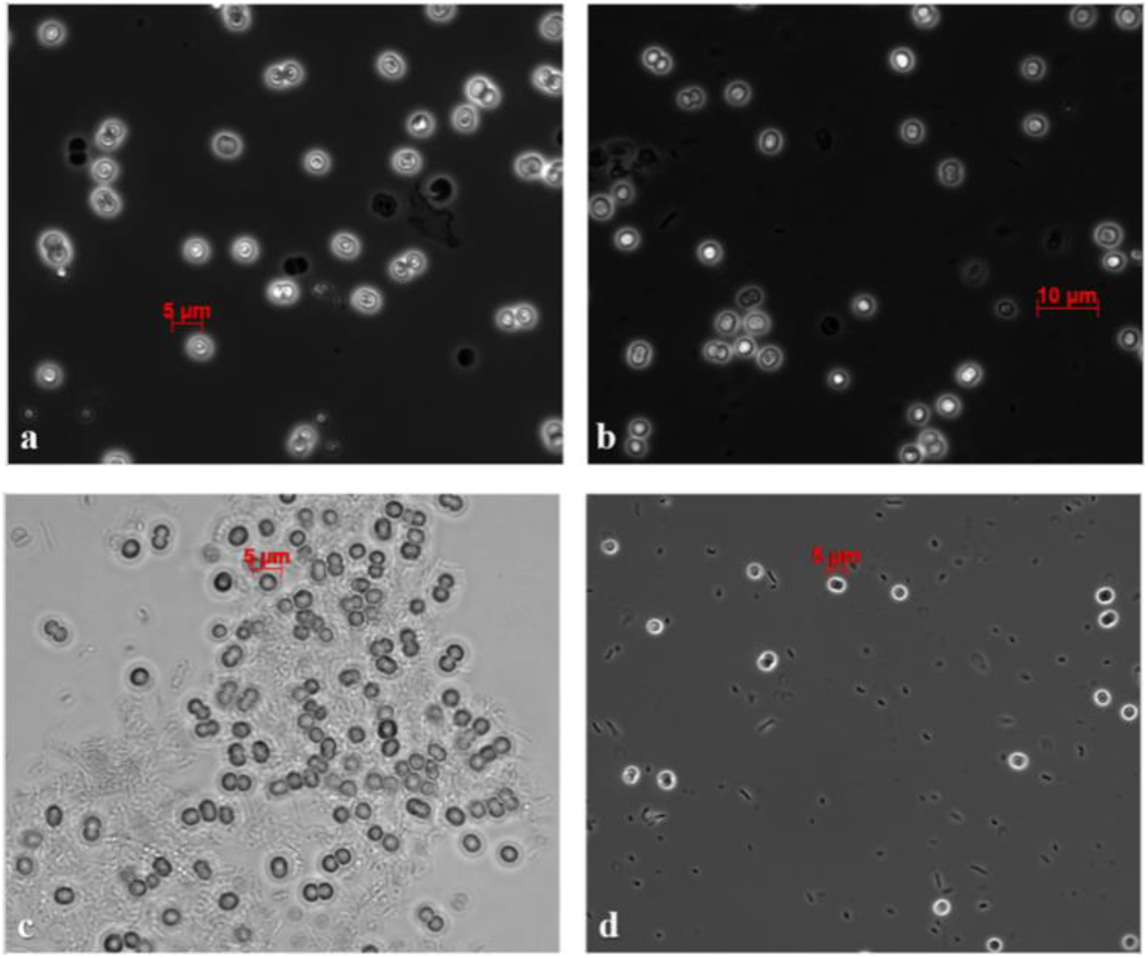
Microscope images from photomixotrophic cultures. Figures *a* – *d* show AxA, LoA, AxA_100_ and LxA_100_ cultures, respectively. Small phototrophic cells (3-4 μm diameter) were identified as well as smaller bacterial cells.

Anaerobic cultures, presented the highest pigment content than aerobic ones. Since the oxygen availability resulted from photosynthesis water oxidation, heterotrophic bacterial growth was disadvantageous. Hence, selection of green phototrophs was possible as some of the phototrophic microorganisms present in the photolitoautotrophic culture (e.g., *Chlorella* sp., *Micractinium* sp., *Scenedesmus* sp., *Desmodesmus* sp., *Synechocystis* sp.) display mixotrophic metabolisms. Some studies demonstrated the possibility of axenic microalgal growth in treated cheese whey (Blier, Lalibert, & de la Notie, 1995; Girard et al., 2014). However, symbiotic relationships in mixed culture are still not explored to their fullest.

As seen in **figure 7**, anaerobic culture on WPC40 and ammonium (CWxA) displayed the highest μ_max_ of all substrates but did not grow as much as cultures in acetate. This can be due to the presence of ordinary heterotrophic organisms together with green phototrophs. The increase in biomass can also be justified by the additional inorganic nutrients for uptake, such as nitrogen and phosphorus that was inherent in CWxA but not in other substrates. Studies on WPC40 in open bottles did not present significant growth (data not shown). Growth rates for different carbon and nitrogen sources and gas exchange allowance are in **table 4**.

**Figure 7.**
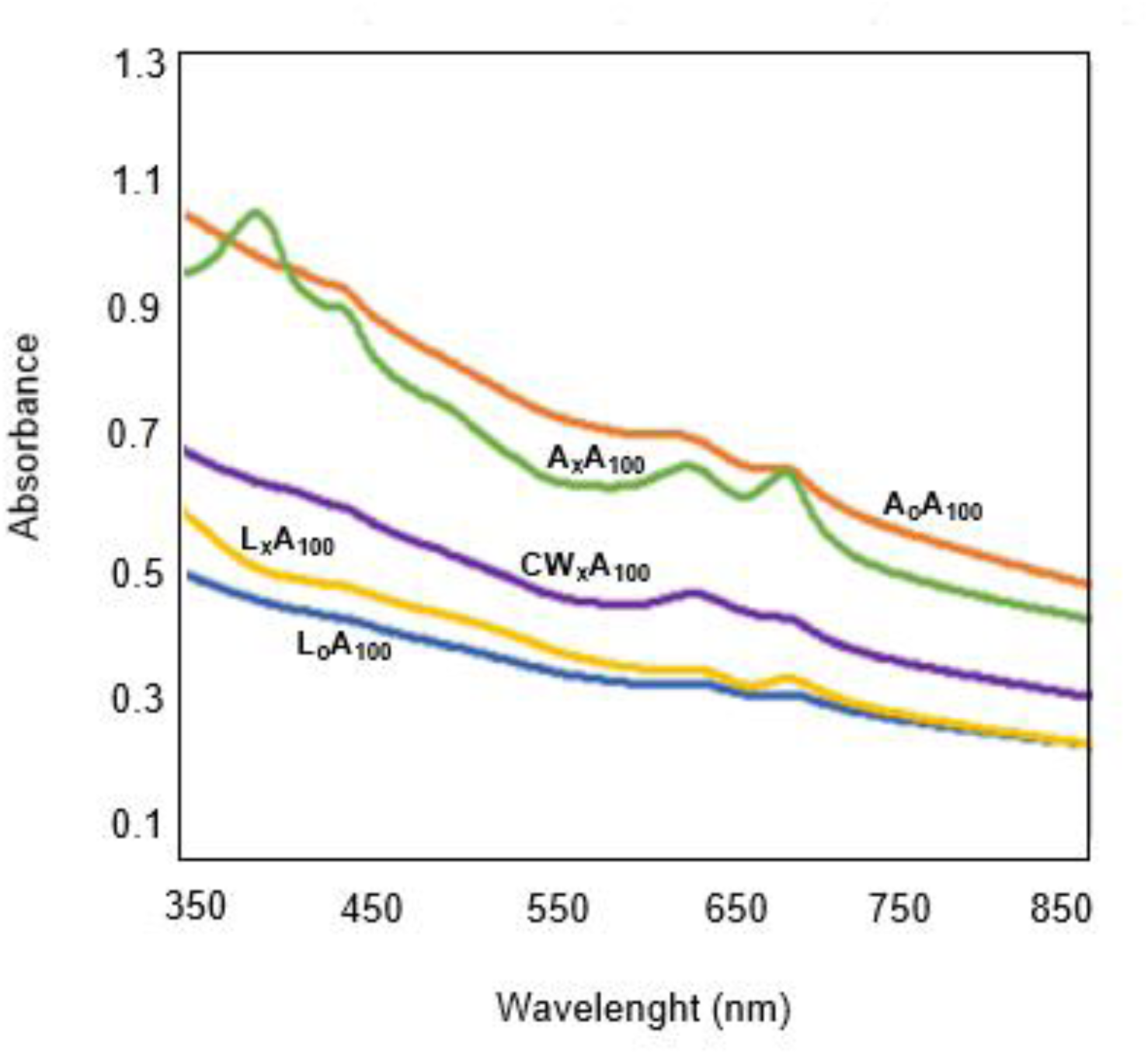
Absorbance spectra curves of A_o_A_100_ (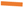), A_x_A_100_ (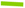), L_o_A_100_ (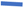), L_x_A_100_ (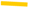) and CWxA_100_ (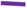) cultures. Peaks related to the presence of pigments are more visible in sealed bottles. Biomass content (A_750_) is similar or higher in open bottles.

**Table 4.** Main characteristics of the cultures grown on ammonia and 18.9 mM-C in aerobic (o) and anaerobic (x) conditions. «+» and «-» signs indicate an increase and a decrease in pH, respectively. “Not available (n.a.)” refers to cultures in which the pigment content was not evaluated.

| Experiment | L<sub>o</sub>A<sub>100</sub> | L<sub>x</sub>A<sub>100</sub> | A<sub>o</sub>A<sub>100</sub> | A<sub>x</sub>A<sub>100</sub> | CW<sub>x</sub>A<sub>100</sub> |
| --- | --- | --- | --- | --- | --- |
| $\mu_{\max}$ (d<sup>-1</sup>) | 0.29 | 0.14 | 0.71 | 0.25 | 0.56 |
| Yield (mg VSS mg-C<sup>-1</sup>) | 0.34 | 0.65 | 0.77 | 1.15 | 0.82 |
| C<sub>max</sub> (mg VSS L<sup>-1</sup>) | 121 | 111 | 189 | 170 | 140 |
| C consumption (%) | 92 | 28 | 96 | 52 | 46 |
| pH (final) | 4.5 | 4.5 | 9.0 | 8.5 | 4.6 |
| Chl<sub>a</sub> (µg mg VSS<sup>-1</sup>) | n.a. | 11.1 | n.a. | 20.3 | 8.6 |
| Chl<sub>b</sub> (µg mg VSS<sup>-1</sup>) | n.a. | 9.3 | n.a. | 13.8 | 9.5 |
| Car (µg mg VSS<sup>-1</sup>) | n.a. | 3.3 | n.a. | 5.3 | 2.3 |

BG-11 medium, a standard medium for the cultivation of cyanobacteria and microalgae, is highly abundant in nitrogen. The C:N:P ratio supplied by the medium is 0.189:101:1. This is significantly imbalanced substrate considering the algal biomass theoretical composition of C_106_H_263_O_110_N_16_P (Redfield, 1934). The excess of nitrogen in this medium is simply used to grow photolithoautotrophs in small batches using atmospheric CO_2_ as additional carbon source.

Carbon, nitrogen, and phosphorus were not a limiting factor in both aerobic and anaerobic cultures. The same could be observed in the C:N:P experiment. However, biomass growth with a carbon concentration of 2.524 mM (C:N:P 36:101:1), showed similar absorbance peaks as the photolitoautotrophic culture. Since nitrogen and phosphorus were a fixed constant, carbon dictated both biomass and pigment concentrations. We can infer that the medium at a C:N:P ratio of 36:101:1, the carbon available was the amount required for biomass and pigment production. The same goes for higher C:N:P ratios. Perhaps, the carbon absorbed was closer to the optimal C:N:P ratio where the remaining carbon was not absorbed by green phototrophs. **Table 5** summarizes the results.

**Table 5.** Variation of the maximum biomass (Bms) growth (A_750_), chlorophyll *a* content (A_680_) and carbon content, and therefore with the C:N ratio.

| C (mmol L<sup>-1</sup>) | C:N:P ratio | Cheese whey |  | Acetate |  |
| --- | --- | --- | --- | --- | --- |
| | | Bms ($A_{750}$) | Chl ($A_{680}$) | Bms ($A_{750}$) | Chl ($A_{680}$) |
| 0.22 | 1:101:1 | 0.29 | 0.37 | 0.83 | 0.92 |
| 0.95 | 5:101:1 | 0.54 | 0.95 | 0.90 | 1.25 |
| 2.52 | 14:101:1 | 0.76 | 1.59 | 0.95 | 1.62 |
| 6.31 | 36:101:1 | 1.2 | 2.33 | 1.13 | 2.37 |
| 12.62 | 72:101:1 | 1.05 | 1.48 | 1.11 | 1.60 |
| 25.24 | 144:101:1 | 1.11 | 1.59 | 0.97 | 1.75 |

Overall, acetate proved to be the most suitable substrate in terms of biomass and pigment concentrations. All chosen organic substrates, WPC40, lactose and acetate, usually have acidic pH. When carbon concentration increased to 18.9 mM C L^−1^, acetate pH increased to 9.0. Acetate cultures in aerobic conditions had a higher pH probably because of atmospheric carbon uptake that allowed a buffer system with some extra carbon availability. In that sense, biomass production could be a result from both photolitoautotrophic and photoorganoheterotrophic metabolisms. As for the acetate in anaerobic conditions, only the photoorganoheterotrophic regime was applied. Since pH was slightly lower than the one on aerobic conditions, we may say that pH was not the limiting factor. The decrease in growth rate in aerobic and anaerobic lactose-ammonium cultures (LoA_100_ and LxA_100_) as well as CWxA_100_ were directly proportional to pH decrease.

So, rather than just pH influence, the suitability of acetate over the other substrates can also be explained by (*i*) efficiency of microorganisms under anaerobic conditions. Besides not having competition with ordinary OHOs, acetate culture under anaerobic conditions microorganisms present a higher growth yield, suggesting that they are able to utilize their resources more efficiently, whereas acetate culture in aerobic conditions microorganisms have a higher growth rate at the rapidly expense of the substrate, (*ii*) the readily uptake of acetate in the glyoxylate cycle. This cycle enables growth on acetate since it bypasses the decarboxylation of the tricarboxylic acid cycle (TAC). Acetate is converted into two molecules of acetyl-CoA by acetyl-CoA synthetase.

Acetyl-CoA can be either a precursor for fatty acids biosynthesis (e.g. chain elongation) or further transformed into succinate for both anabolic and catabolic processes.

### 3.3 Transition of metabolic regimes and substrates in a SBR-CSTR

#### 3.3.1 Photolitoautotrophic acclimation for selection for green phototrophs under batch mode

A stirred-tank photobioreactor was initially operated in batch mode for acclimation of photolitoautotrophic culture prior to transition of substrate and metabolic regime. During the first six days, a similar growth rate (0.99 d^−1^) to the one with undissolved carbonate flasks was observed. When carbonate minerals were fully dissolved growth was limited, contrary to what was observed in the undissolved carbonate flasks. This can be due to complete carbon consumption in BG-11 medium and the inability of atmospheric carbon uptake giving the anaerobic mode. The fact that the gas flow consisted of argon, limited any possibility of carbon source or inorganic nutrients entry in the system.

Nitrogen and phosphorus measurements, (>130 mg N-NO_3_^−^ L^−1^ and > 5 mg P-PO_4_^3-^) confirmed that the nutrients available in BG-11 were sufficient to sustain microalgae growth. Giving the initial high substrate concentration in batch regimes, microorganisms were selected according growth rate. **Figure 8** compares photolitoautotrophic biomass growth in different batch scales.

**Figure 8.**
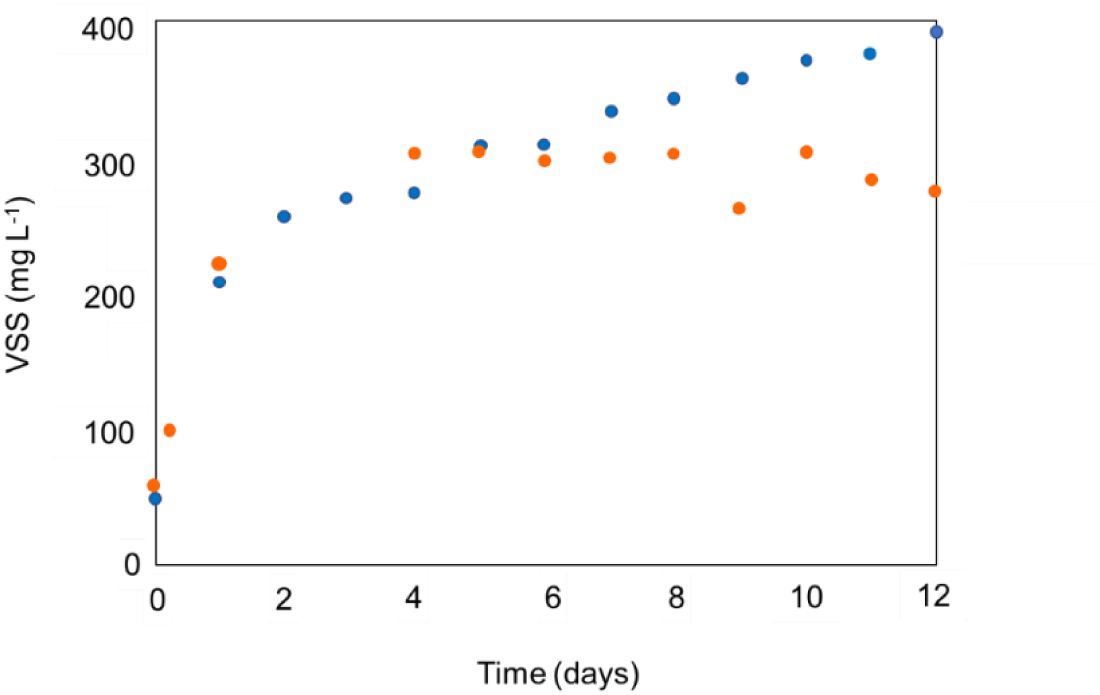
Biomass growth in shake-flask culture (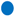) and photobioreactor – batch mode (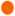). Growth in the photobioreactor was limited after six days due to atmospheric carbon dioxide dissolution limitation.

#### 3.3.2 Lower dilution rate favors green phototrophs selection in a chemostat

Acetate replace BG-11 on the transition from photolitoautotrophic regime to a photoorganoheterotrophic one. An initial dilution rate of 0.24 d^−1^ displayed a biomass concentration of 41 mg VSS d^−1^. The selection of this specific dilution rate was a result of the maximum growth rate of 0.46 d^−1^ observed in the aerobic acetate-ammonium and 0.189 mM C L^−1^, as in BG-11. However, at this dilution rate the culture had a few phototrophic cells which can be seen in the in the absorbance spectra of the sample, especially on the wavelengths corresponding to pigments (440, 630 and 680 nm). **Figure 9** shows the morphologic characterization of microorganisms community.

**Figure 9.**
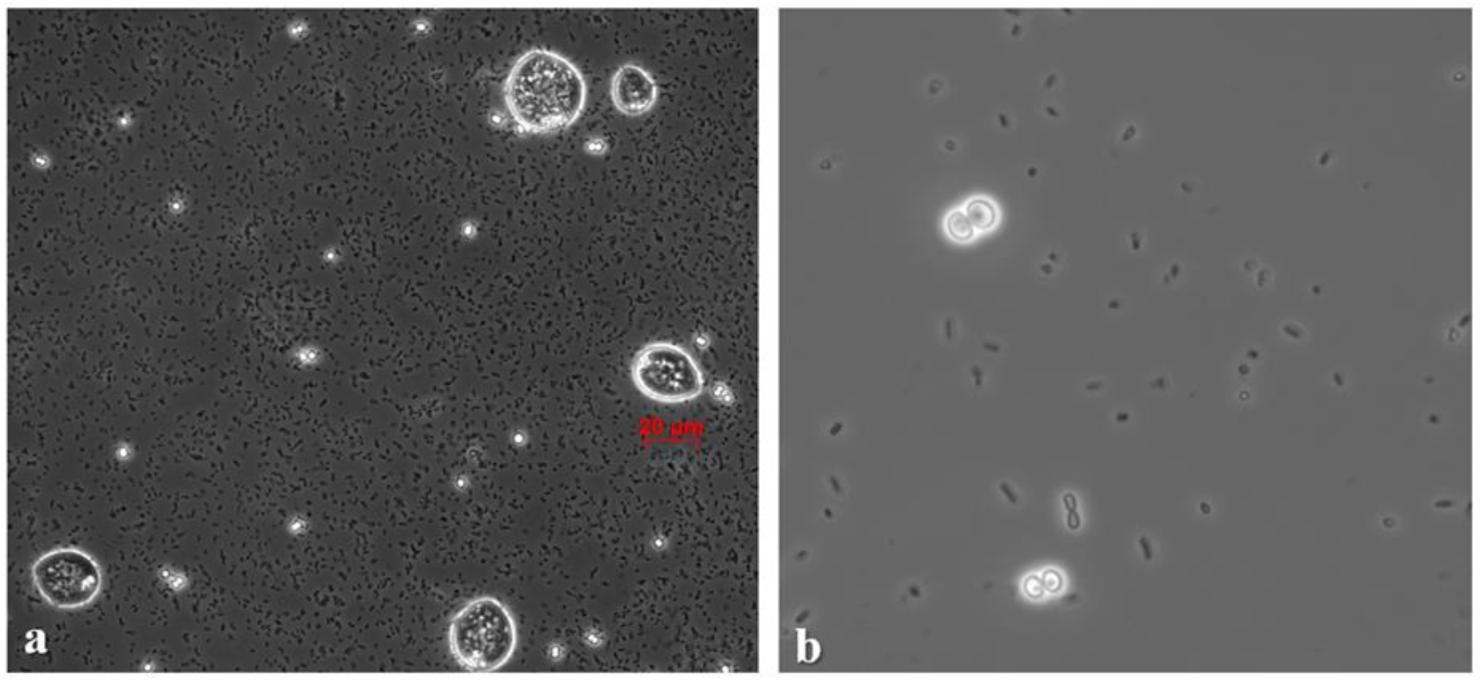
Microscope images of the photomixotrophic culture cultivated on acetate in the CSTR PBR. Image ***a*** was taken from the cultivation with a dilution rate of 0.25 d^−1^ and image ***b*** from the 0.17 d^−1^ dilution rate cultivation.

A different dilution rate of 0.17 d^−1^ was then selected based on the maximum growth rate of 0.25 d^−1^ observed anaerobic acetate-ammonium and 18.9 mmol C L^−1^ as the experiment in the previous section. The dilution rate alteration led to visible improvements as reactor presented a dark green content, an improved absorbance spectrum and an increase in biomass concentration from 86 to 125 mg VSS L^−1^. Presence of ciliates and smaller bacteria was still visible in the culture although in less quantity than phototrophic cells as observed in **figure 9 a**.

The dissolved oxygen in the culture increased from 0% to 2%. The carbon dioxide in the off-gas decreased from 0.085% to 0.058%, and the transition was oscillatory, indicating a positive symbiotic relationship with adequate retention time. A picture of the CSTR, spectra profile from batch mode and both dilution rates and biomass evolution can be seen on **figure 10**. **Table 6** depicts the CSTR growth parameters in both dilution rates.

**Figure 10.**
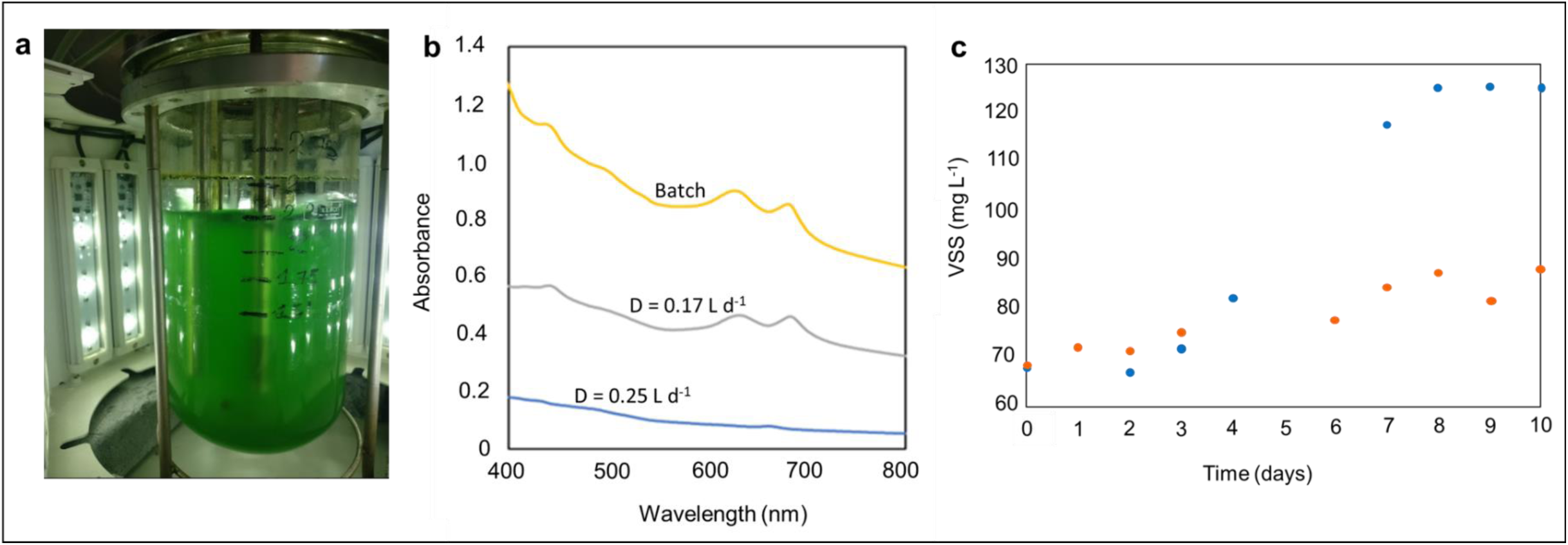
a. CSTR PBR with a dilution rate of 0.17 d^−1^, **b** Absorbance spectra of the microalgal cultivation on acetate in a CSTR with 0.17 d^−1^ and 0.25 d^−1^ dilution rates and in a batch (AxA_b100_). Higher dilution rate caused phototrophic biomass wash out. At a lower dilution rate, absorbance peaks were identified at the same wavelengths as in the batch culture (AxA_b100_). **c** Biomass growth at different dilution rates (0.25 d^−1^ and 0.17 d^−1^). The 0.17 d^−1^ dilution rate allowed higher biomass growth inside the reactor. Dilution rate was the only different parameter between both experiments.

**Table 6.** Growth parameters of the mixed culture grown on acetate (18.9 mmol C L^−1^), using ammonium as a N-source, in CSTR mode for two different dilution rates.

| Dilution rate ($\text{d}^{-1}$) | Yield<br>($\text{mg VSS mg C}^{-1}$) | Carbon removal<br>(%) | $\text{C}_{\text{vss}}$<br>($\text{mg VSS L}^{-1}$) | $\text{P}_{\text{vss}}$<br>($\text{mg VSS d}^{-1}$) | $\text{P}_{\text{CO}_2}$<br>($\text{mL-CO}_2 \text{ d}^{-1}$) | $\text{q}_s$<br>($\text{mg}_{\text{acetate}} \text{ mg VSS}^{-1} \text{ d}^{-1}$) |
| --- | --- | --- | --- | --- | --- | --- |
| **0.24** | 0.45 | 93 | 86 | 41 | 62.6 | 1.30 |
| **0.17** | 0.66 | 95 | 125 | 45 | 39.6 | 1.38 |

## 4 Conclusions

This work aimed on the ecologic engineering of green photoorganoheterotrophic microalgal biomass growth for cheese whey full valorization. For that, we suggested the couple microalgae biomass growth process with cheese whey acidogenic fermentation processes given their potential 66.5 million m^3^ y^−1^ production destined to lower aggregated-value products or depending on location, its discharge into water bodies leading to eutrophication. The following can be concluded:

1. Green phototrophs selection from channel surface water was successful and pigment content was directly proportional to biomass concentration.
2. Despite changing the nitrogen source from nitrate to ammonium, denitrifiers were able to metabolize both nitrogen sources and use the dissolved oxygen in the medium as electron acceptor.
3. In batch experiments, acetate was the most suitable organic carbon source for biomass and pigments production with a maximum growth rate of 0.25 d^−1^ and biomass concentration of 170 mg VSS d^−1^, compared to lactose (0.29 d^−1^ and 111 mg VSS d^−1^), and WPC40 (0.56 d^−1^ and 140 mg VSS d^−1^). The higher growth rate presented in WPC40 can be a result of additional inorganic nutrients.
4. The uptake of acetate was is thought to take place via the glyoxylate cycle for chain elongation, given the absence of other carbon sources in the CSTR reactor.
5. Selection of green phototrophs were successful under chemostat regime with a lower dilution rate to prevent microalgae wash-out and out-competition with OHOs, displaying 125 mg L^−1^ of biomass concentration and 45 mg VSS d^−1^ of biomass production.
6. The use of argon as the off-gas carrier together with the presence of dissolved oxygen suggests a symbiotic relationship within the microbial community.

Results on the amplicon gene sequencing of batches and both chemostat dilutions are currently being assessed and will shed light on the behavior of green photoorganoheterotrophs in mixed cultures. In addition, acetate metabolism of potoorganoheterotrophs should be better understood in order to diverge their metabolism to the glyoxylate cycle and optimize the process for both upscaling and cheese whey full valorization.

